## Supplementary material for "Autophagy across tissues of aging mice": Figure S

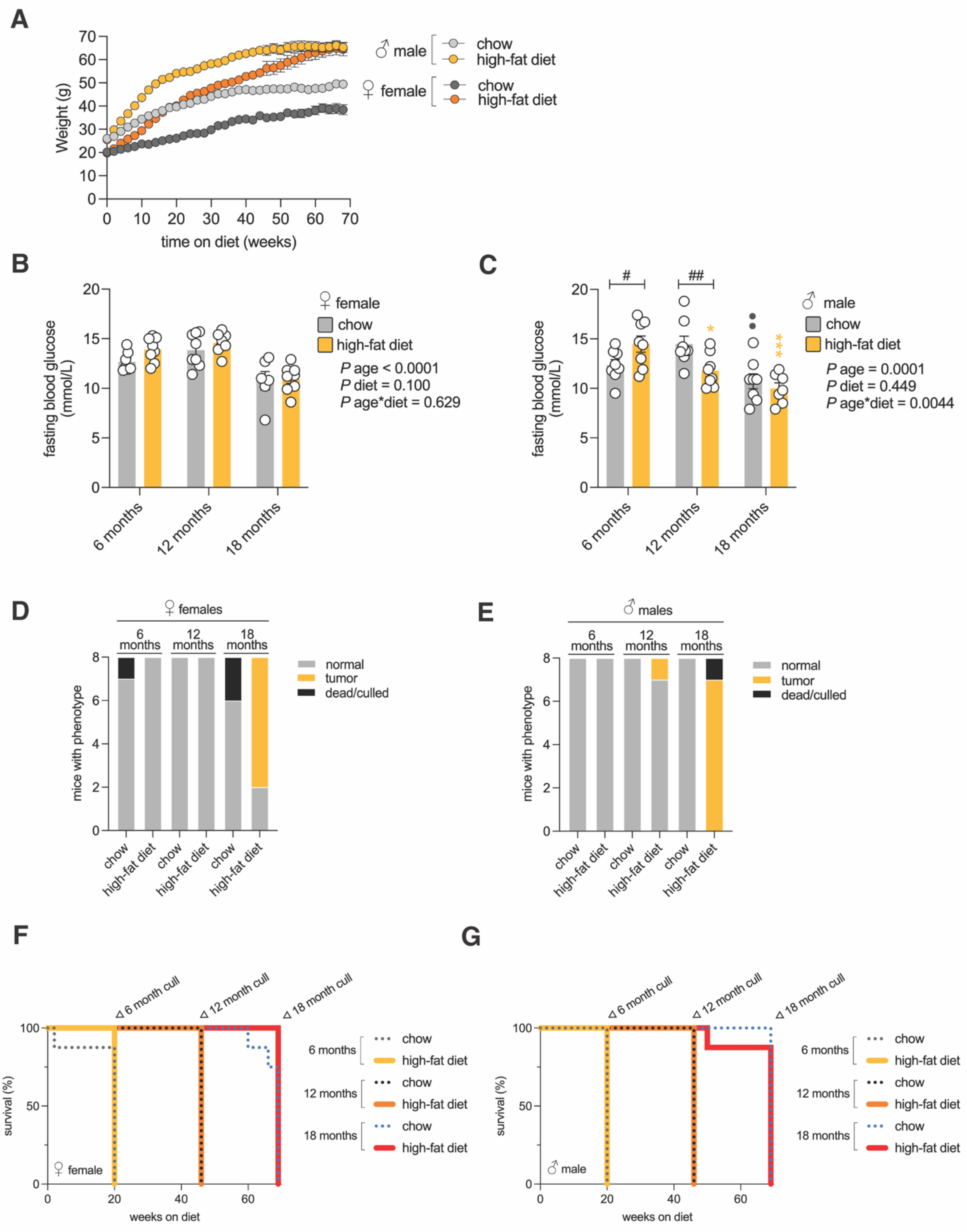


**Figure S1. Metabolic and physiological effects of aging and high-fat diet.**

A. Body weight of tf-LC3B mice that were fed regular chow or a high-fat diet**.** Values are mean body weight (g) ± sem for n = 6-24 mice/sex/diet/timepoint.

B. Fasting blood glucose levels of female tf-LC3B mice that were fed regular chow or a high-fat diet**.** Values are blood mean glucose levels (mmol/L) ± sem for n = 6-8 female mice/diet/timepoint (2-way ANOVA).

C. Fasting blood glucose levels of male tf-LC3B mice that were fed regular chow or a high-fat diet**.** Values are mean blood glucose levels (mmol/L) ± sem for n = 7-8 male mice/diet/timepoint (2-way ANOVA with Tukey’s multiple comparisons test).

D. Tumor incidence and mortality rates of female tf-LC3B mice that were fed regular chow or a high-fat diet; n values (mice/sex/diet/timepoint) are indicated by phenotype.

E. Tumor incidence and mortality rates of male tf-LC3B mice that were fed regular chow or a high-fat diet; n values (mice/sex/diet/timepoint) are indicated by phenotype.

F. Survival analysis of female tf-LC3B mice that were fed regular chow or a high-fat diet. n = 8 female mice/diet/timepoint.

G. Survival analysis of male tf-LC3B mice that were fed regular chow or a high-fat diet. n = 8 female mice/diet/timepoint.

Statistically significant p values comparing diet effects (#); age effects within each diet (coloured) compared to 6-month time point (*) or comparing 12- and 18-month time points (●).

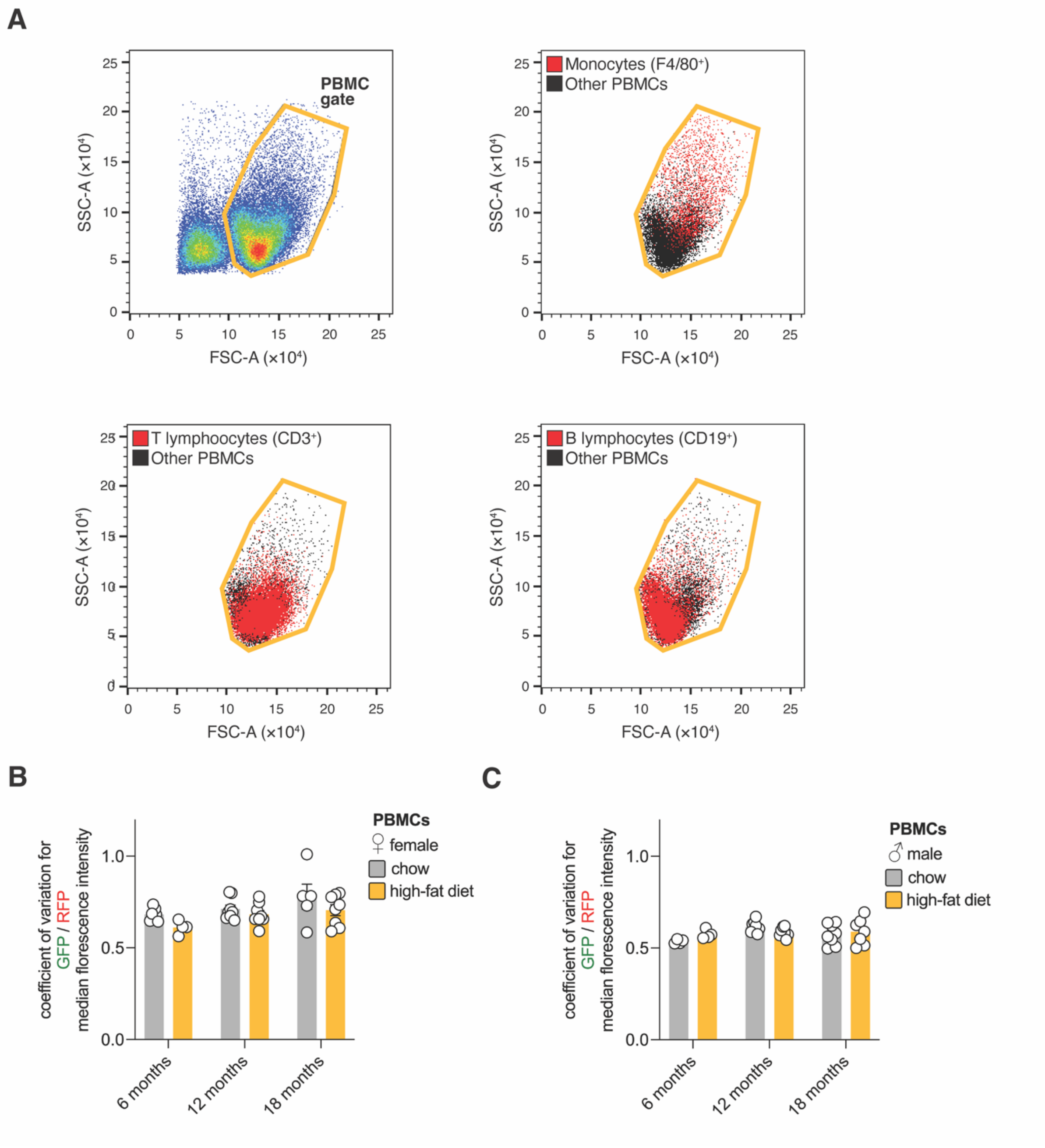


**Figure S2. Verification of PBMCs.**

A. PBMCs purified from blood are enriched with monocytes as well as T and B lymphocytes. PBMCs extracted from C57BL/6J (non-transgenic) mice were immunostained for monocyte (F4/80) as well as T- (CD3) and B-lymphocyte (CD19) markers and analyzed via flow cytometry. This illustrates that gating parameters employed were appropriate.

B. Variation of autophagic flux in female PBMCs. Values are coefficient of variation in GFP/RFP median fluorescence intensity across cells within each animal. Data is related to Figure 2A.

C. Variation of autophagic flux in male PBMCs. Values are coefficient of variation in GFP/RFP median fluorescence intensity across cells within each animal is shown. Data is related to Figure 2B.

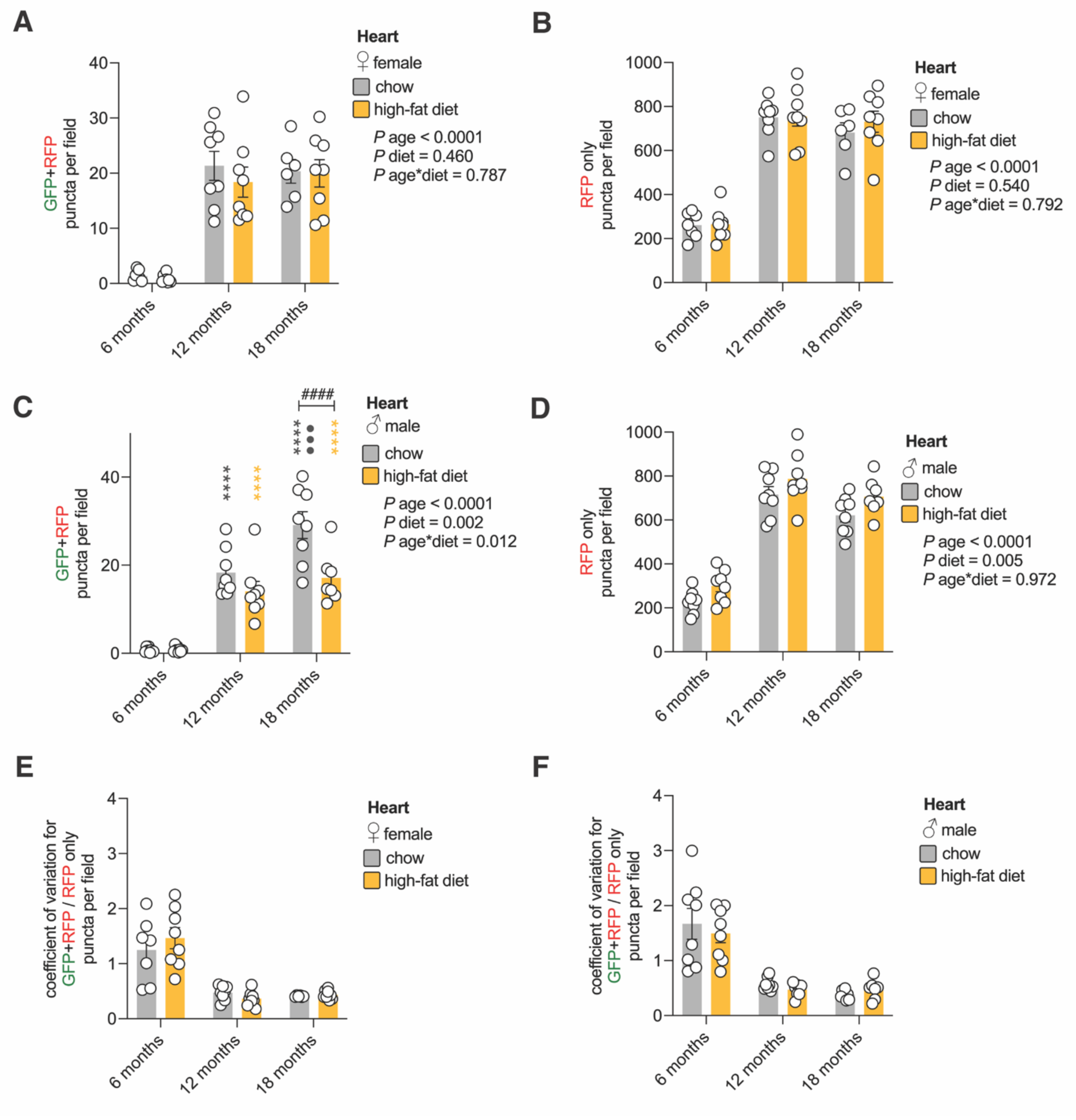


**Figure S3. Autophagosome and autolysosome abundance, and autophagic flux variability in the heart.**

A. Quantification of autophagosomes in the female heart. Values are GFP+RFP puncta per field ± sem for n = 6-8 female mice/timepoint (2-way ANOVA).

B. Quantification of autolysosomes in the female heart. Values are RFP only puncta per field ± sem for n = 6-8 female mice/timepoint (2-way ANOVA).

C. Quantification of autophagosomes in the male heart. Values are GFP+RFP puncta per field ± sem for n = 7-8 male mice/timepoint (2-way ANOVA with Tukey’s multiple comparisons test).

D. Quantification of autolysosomes in the male heart. Values are RFP only puncta per field ± sem for n = 7-8 male mice/timepoint (2-way ANOVA).

E. Variation of autophagic flux in the female heart. Coefficient of variation in GFP+RFP/RFP only puncta per field across 10 fields for each animal. Data is related to Figure 3A and B.

F. Variation of autophagic flux in the male heart. Coefficient of variation in GFP+RFP/RFP only puncta per field across 10 fields for each animal. Data is related to Figure 3C and D.

Statistically significant p values comparing diet effects (#); age effects within each diet (coloured) compared to 6-month time point (*) or comparing 12- and 18-month time points (•).

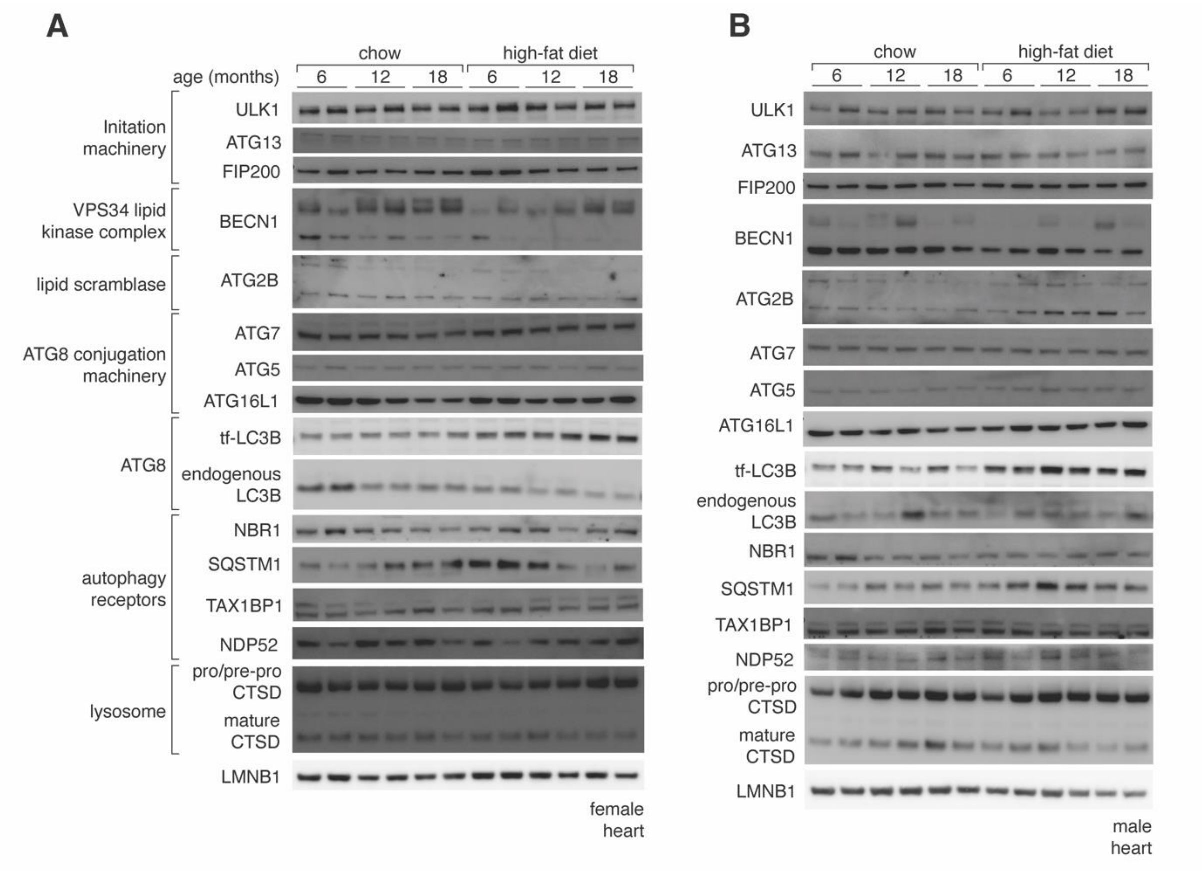


**Figure S4. Abundance of key autophagy proteins in the heart.**

A. Immunoblot analysis of heart lysates from female tf-LC3B mice that were fed regular chow or a high-fat diet**.** Blots were probed as indicated. n = 2 mice/diet/timepoint is shown.

B. Immunoblot analysis of heart lysates from male tf-LC3B mice that were fed regular chow or a high-fat diet**.** Blots were probed as indicated. n = 2 mice/diet/timepoint is shown.

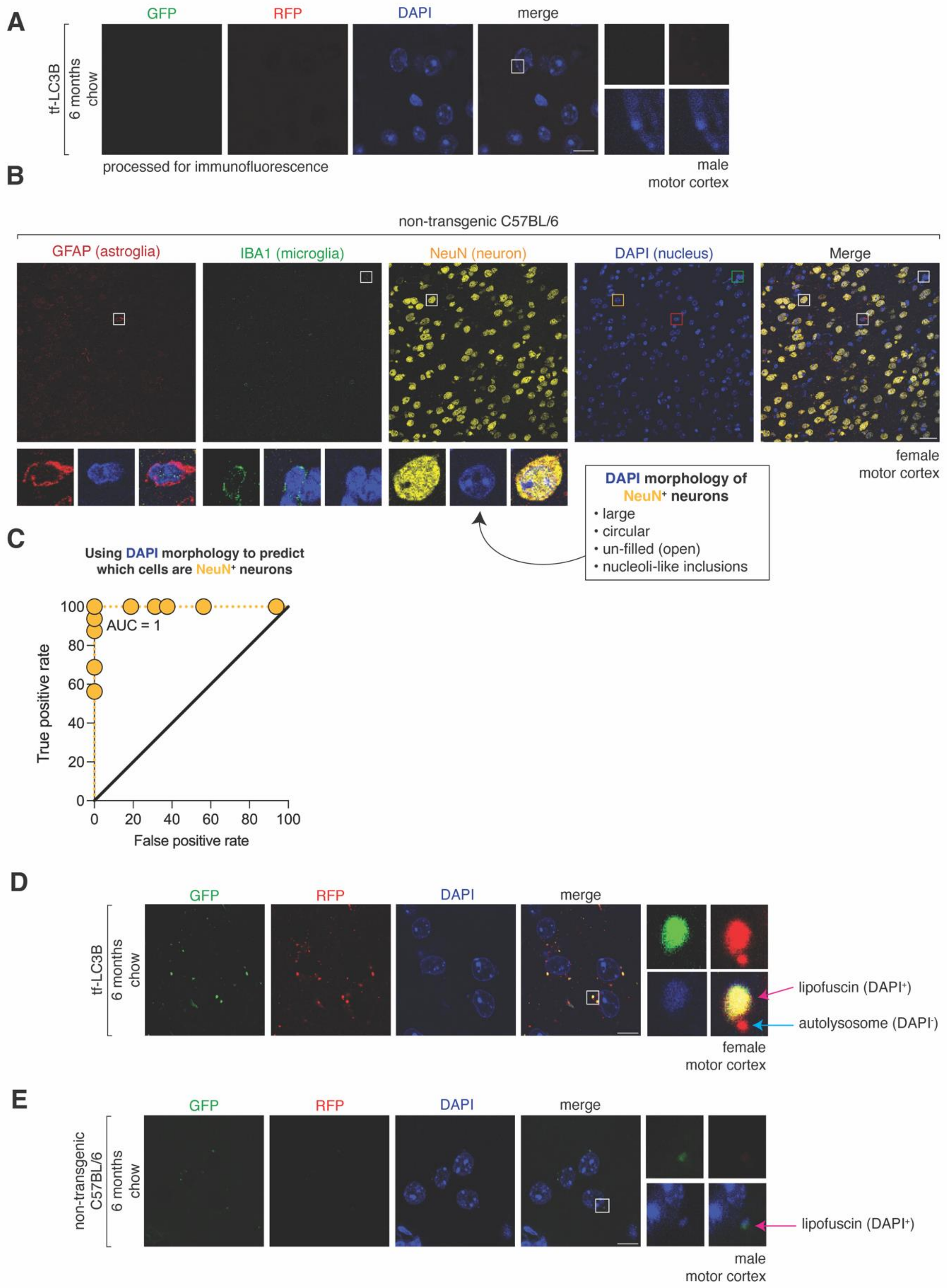


**Figure S5. Large DAPI nuclei as a proxy for identifying NeuN$^{+}$ neurons, and DAPI puncta for detection of lipofuscin in the brain.**

A. Immunofluorescence processing quenches GFP and RFP signals. Confocal images of the brain of tf-LC3B mice after processing for immunofluorescence (without primary or secondary antibodies). Scale bar: 10 μm. Inset: 10× magnification. Blue on merge: DAPI.

B. DAPI-stained nuclei morphology of $NeuN^+$ neurons. Confocal images of the motor cortex of female non-transgenic control mice that were fed regular chow for 12-months, immunostained for neuronal (NeuN), astroglial (GFAP), and microglial (IBA1) markers. Scale bar: 30 μm. Inset: 8× magnification. Blue on merge: DAPI.

C. Morphological features of DAPI-stained nuclei (large size, circular shape, unfilled/ “open” lumen, and nucleoli-like inclusions) can reliably distinguish $NeuN^+$ neurons when used as selection criteria by a blinded experimenter. Approximately 1400 $DAPI^+$ nuclei/cells from n = 2 mice/sex were analyzed (receiver operating characteristic curve analysis; AUC = 1).

D. Identification of lipofuscin autofluorescence in the brain. Confocal images of the brain of tf-LC3B mice. The image shown is duplicated from (Figure 4A) but highlights evidence of fluorescent artefacts caused by lipofuscin in the inset. Scale bar: 10 μm. Inset: 10× magnification. Blue on merge: DAPI.

E. Identification of lipofuscin autofluorescence in the brain. Confocal images of the brain of non-transgenic C57BL/6 mice. Fluorescent artefacts caused by lipofuscin are displayed in the inset. Scale bar: 10 μm. Inset: 10× magnification. Blue on merge: DAPI.

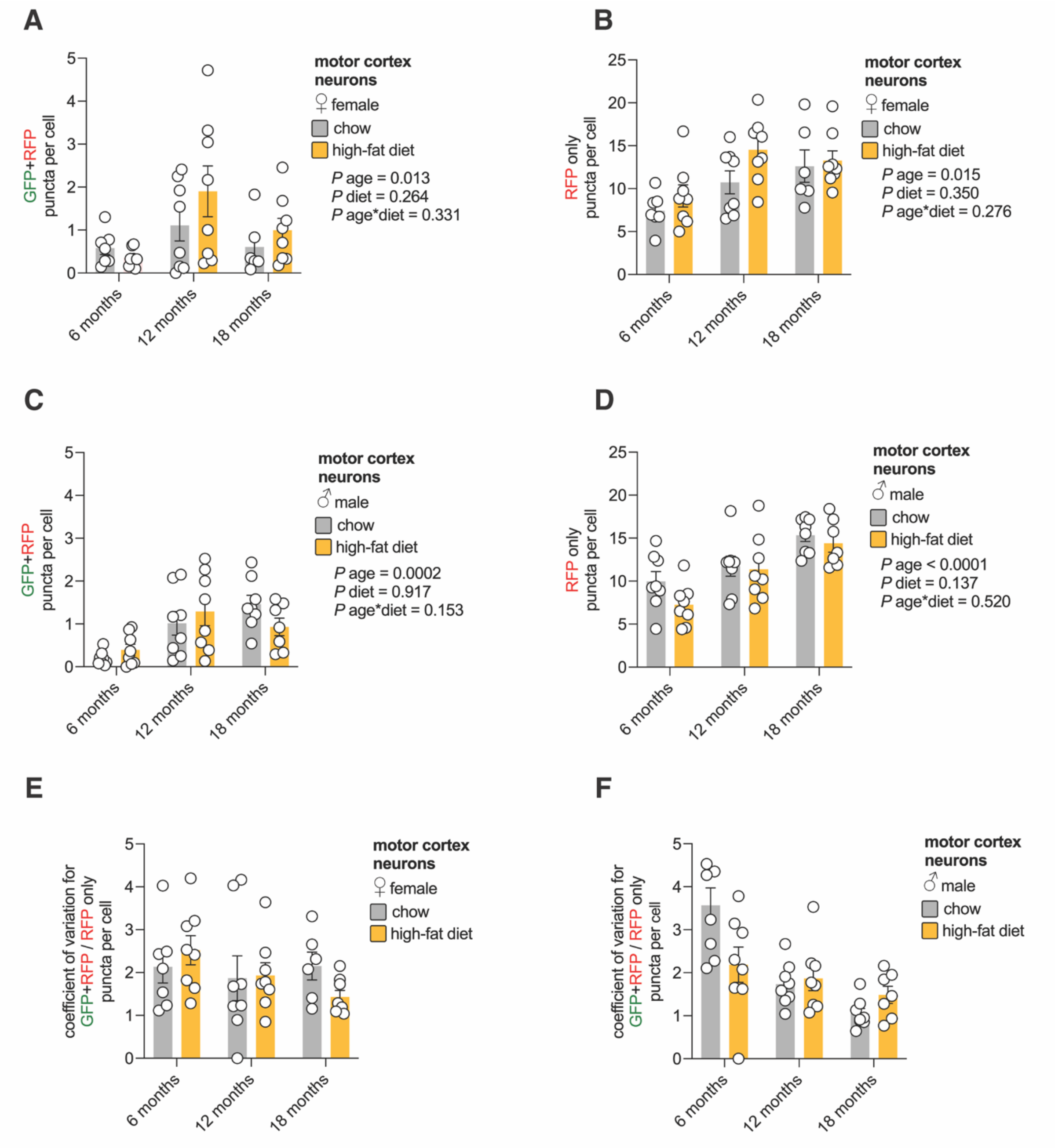


**Figure S6. Autophagosome and autolysosome abundance, and autophagic flux variability in motor cortex neurons.**

A. Quantification of autophagosomes in female motor cortex neurons. Values are GFP+RFP puncta per field ± sem for n = 6-8 female mice/timepoint (2-way ANOVA).

B. Quantification of autolysosomes in female motor cortex neurons. Values are RFP only puncta per field ± sem for n = 6-8 female mice/timepoint (2-way ANOVA).

C. Quantification of autophagosomes in male motor cortex neurons. Values are GFP+RFP puncta per field ± sem for n = 7-8 male mice/timepoint (2-way ANOVA).

D. Quantification of autolysosomes in male motor cortex neurons. Values are RFP only puncta per field ± sem for n = 7-8 male mice/timepoint (2-way ANOVA).

E. Variation of autophagic flux in female motor cortex neurons. Coefficient of variation in GFP+RFP/RFP only puncta per cell across 10 fields for each animal. Data is related to Figure 4A and B.

F. Variation of autophagic flux in male motor cortex neurons. Coefficient of variation in GFP+RFP/RFP only puncta per cell across 10 fields for each animal is shown. Data is related to Figure 4C and D.

Statistically significant p values comparing diet effects (#); age effects within each diet (coloured) compared to 6-month time point (*) or comparing 12- and 18-month time points (•).

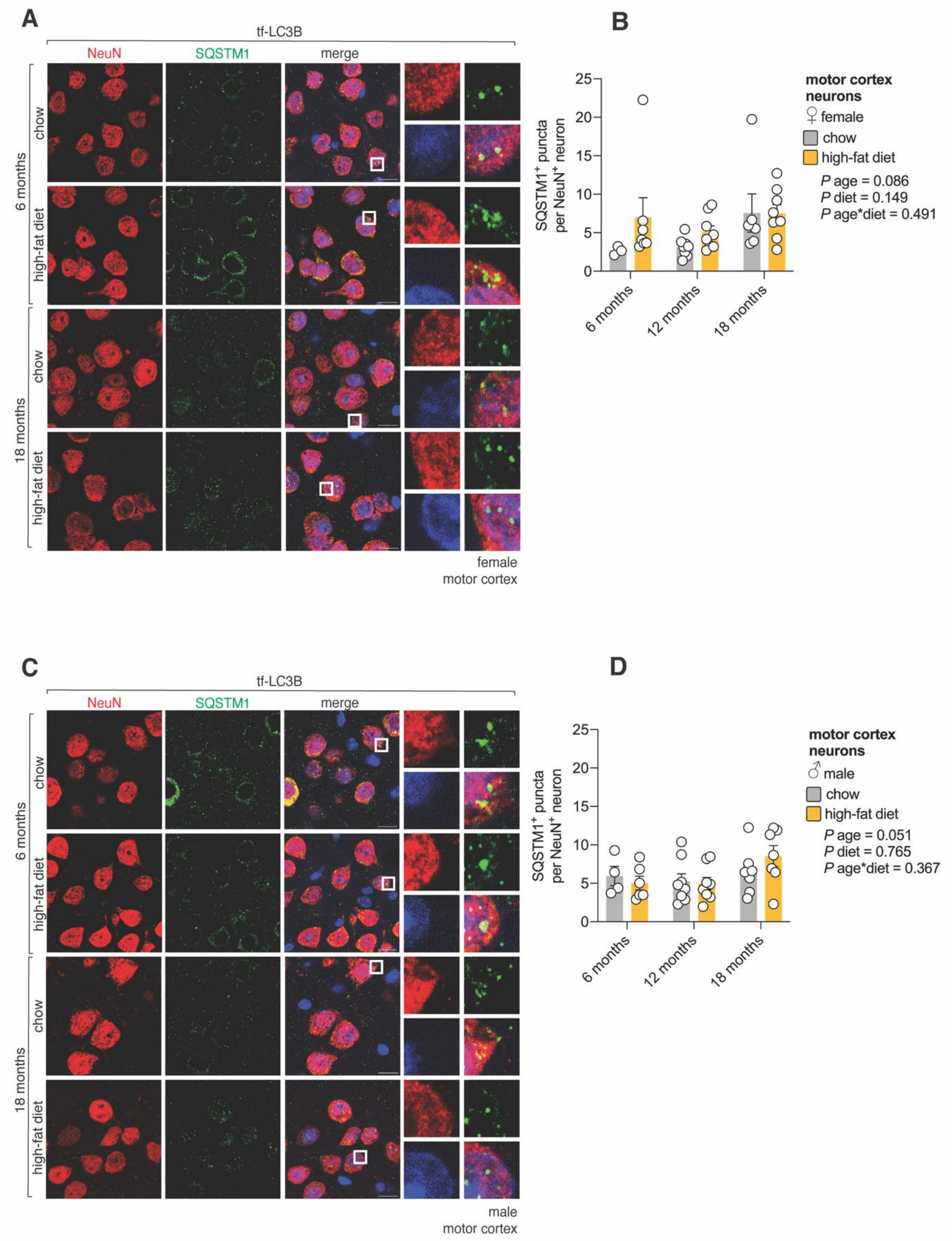


**Figure S7. SQSTM1/p62 inclusions within motor cortex neurons.**

A. SQSTM1/p62 inclusions in female motor cortex neurons. Confocal images of the brain of female tf-LC3B mice that were fed regular chow or a high-fat diet for 6- and 18-months (12-month timepoint not shown), immunostained for SQSTM1/p62 and the neuronal marker NeuN.

Note that our immunofluorescence protocol quenched GFP and RFP signals from tf-LC3B (Figure S3A) allowing those channels to be used for detection. Scale bar: 10 $\mu$m. Inset: 10$\times$ magnification. Blue on merge: DAPI

B. Quantification of SQSTM1/p62 inclusions in female brain neurons. Values are SQSTM1$^+$ puncta per NeuN$^+$ neuron for n = 3-8 female mice/timepoint (2-way ANOVA).

C. SQSTM1/p62 inclusions in male brain neurons. Confocal images of the brain of male tf-LC3B mice that were fed regular chow or a high-fat diet for 6- and 18-months (12-month timepoint not shown), immunostained for SQSTM1/p62 and the neuronal marker NeuN. Scale bar: 10 $\mu$m. Inset: 10$\times$ magnification. Blue on merge: DAPI.

D. Quantification of SQSTM1/p62 inclusions in male brain neurons. Values are SQSTM1$^+$ puncta per NeuN$^+$ neuron for n = 4-8 male mice/timepoint (2-way ANOVA).

Statistically significant p values comparing diet-effects (#); age-effects within each diet (coloured) compared to 6-month time point (*) or comparing 12- and 18-month time points (•).

female heart

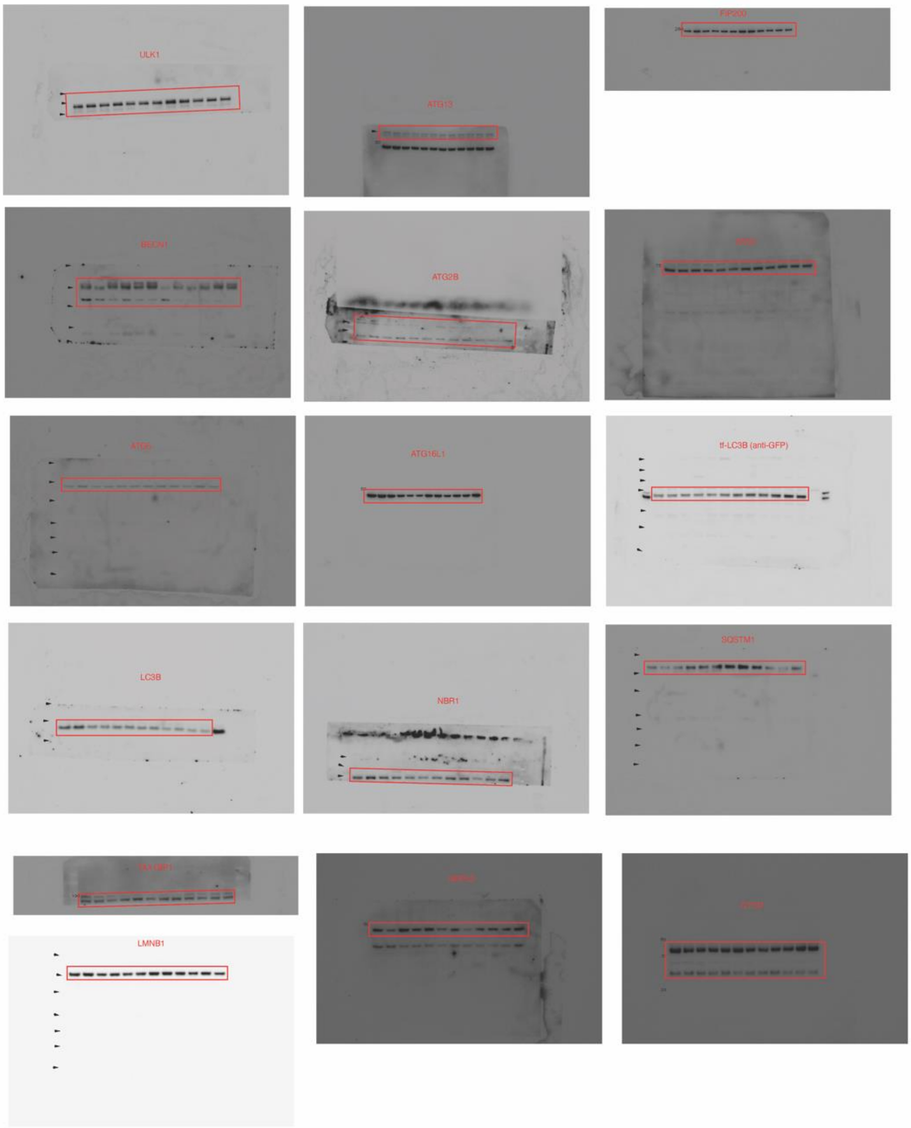

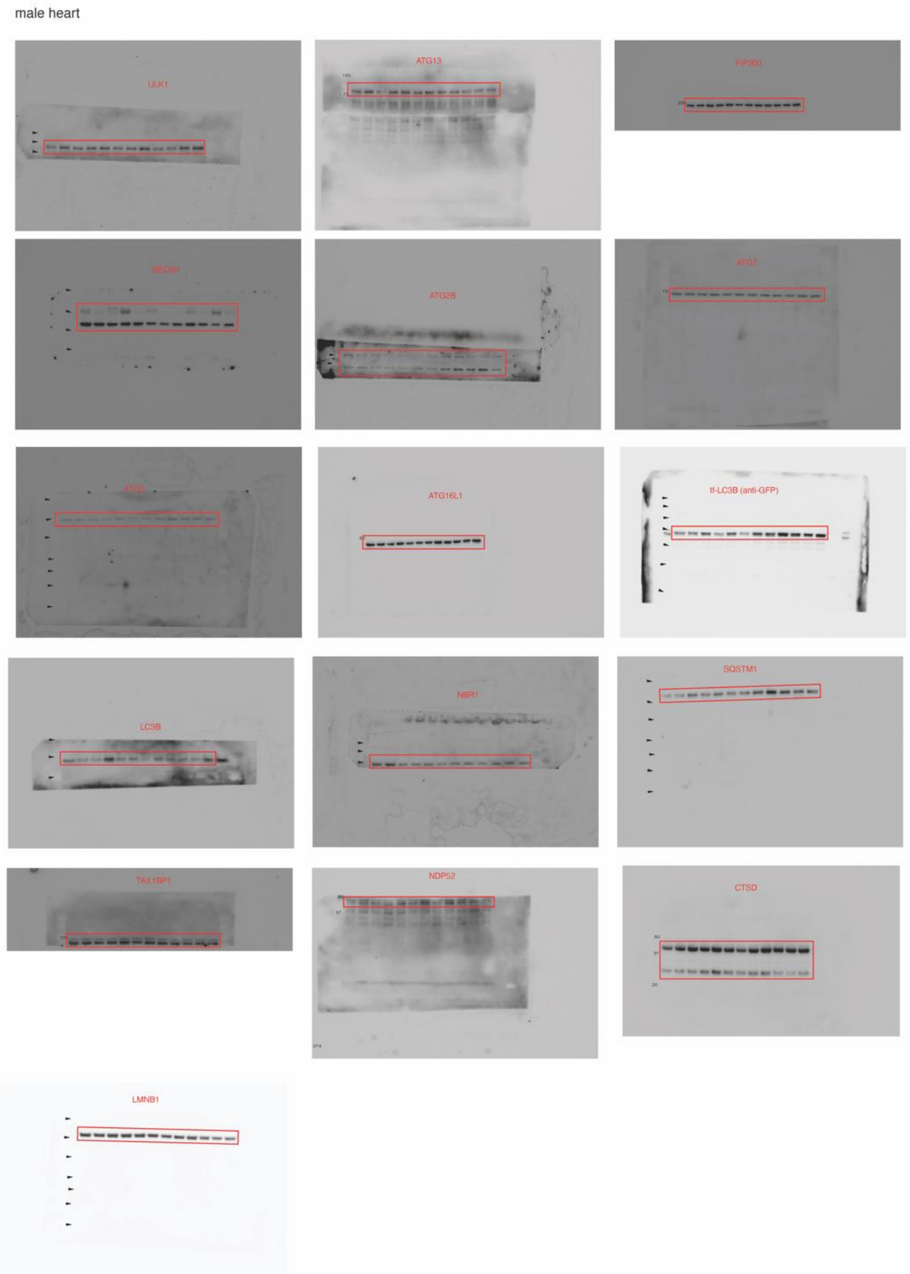


**Figure S8. Uncropped immunoblots from Figure S4.**
